# BugSplit: highly accurate taxonomic binning of metagenomic assemblies enables genome-resolved metagenomics

**DOI:** 10.1101/2021.10.16.464647

**Authors:** Induja Chandrakumar, Nick P.G. Gauthier, Cassidy Nelson, Michael B. Bonsall, Kerstin Locher, Marthe Charles, Clayton MacDonald, Mel Krajden, Amee R. Manges, Samuel D. Chorlton

## Abstract

A large gap remains between sequencing a microbial community and characterizing all of the organisms inside of it. Here we develop a novel method to taxonomically bin metagenomic assemblies through alignment of contigs against a reference database. We show that this workflow, BugSplit, bins metagenome-assembled contigs to species with a 33% absolute improvement in F1-score when compared to alternative tools. We perform nanopore mNGS on patients with COVID-19, and using a reference database predating COVID-19, demonstrate that BugSplit’s taxonomic binning enables sensitive and specific detection of a novel coronavirus not possible with other approaches. When applied to nanopore mNGS data from cases of *Klebsiella pneumoniae* and *Neisseria gonorrhoeae* infection, BugSplit’s taxonomic binning accurately separates pathogen sequences from those of the host and microbiota, and unlocks the possibility of sequence typing, *in silico* serotyping, and antimicrobial resistance prediction of each organism within a sample. BugSplit is available at https://bugseq.com/academic.

## Introduction

Automated genome-resolved metagenomics, including the identification and characterization of members of a microbial community, has remained challenging despite improvements in sequencing technology and bioinformatic analysis^1^. A community-initiative for the Critical Assessment of Metagenomics Interpretation (CAMI) has tracked the progress of this goal over time and posited two main challenges for metagenomic next-generation sequencing (mNGS): profiling and binning^2^. Metagenomic profiling aims to quantify the presence/absence and abundance of organisms in a microbial community, and has seen a marked improvement in the number and performance of tools from the first to the second CAMI challenge^3^.

Metagenomic binning aims to place sequences (usually assembled contigs) from the same organism in a unique bin, enabling the study of each organism within complex microbial communities. Metagenomic binning can further be divided into supervised and unsupervised approaches, where unsupervised approaches use sequence information such as tetranucleotide repeat counts and sequencing depth to bin sequences, while supervised approaches use reference sequence databases and previously generated information to bin sequences^4^. While unsupervised binners have improved over recent years, their lack of ability to assign taxonomic labels to bins precludes downstream analyses such as profiling the presence/absence of specific organisms, performing organism-specific analyses, or identifying sequences of concern (eg. novel pathogens with sequence homology to known pathogens)^3^. The COVID-19 pandemic has highlighted the need for such improvements, with the aim of ensuring early availability of pathogen-agnostic diagnostics when outbreaks of novel strains emerge^5^.

Earlier work on taxonomic binning has relied on amino acid alignments of assembled contigs to a universal protein database^6–9^. These workflows allow for identification of divergent sequences, but do not leverage the non-coding and synonymous variation within contigs, nor the positional relationship of classifier features (eg. co-localization of proteins) into taxonomic classification. An alternative approach relies on a search for taxa-specific conserved features; however, this approach is limited by recall, where contigs missing the conserved marker cannot be classified, and to date, no approaches encompass conserved markers spanning archaea, bacterial, viruses and eukaryotes^10,11^. Finally, k-mer and minimizer based approaches suffer from lack of positional relationship between k-mers, and lack of ability to resolve uncertainty when using a single k-mer or even base to break lowest common ancestor ties, as demonstrated in previous evaluations^9^.

In addition to software improvements, advancements in sequencing technology have enabled the production of higher quality metagenomic assemblies through long-read approaches, which have been posited to enable genome-resolved metagenomics^12,13^. Third-generation sequencers, such as those by Oxford Nanopore Technologies (ONT) and Pacific Biosciences (PacBio), can generate reads up to megabases in length and contigs spanning full microbial genomes^14^. Traditionally, there has been a tradeoff between read length and sequencing quality; however with the advent of high fidelity (HiFi) PacBio sequencing and improvements in ONT sequencing kits and basecallers, that gap is quickly disappearing^15–17^. Additionally, the small size and costeffectiveness of third-generation sequencers is poised to enable broad-scale adoption of this technology for mNGS, which has already been used across clinical, food, defense and environmental applications^18–20^.

Here we describe BugSplit, an automated, supervised method to bin contigs by taxonomic identity. The main difference between our workflow, BugSplit, and others is that it utilizes local nucleotide alignments of contigs against a universal reference database, such as the NCBI nucleotide (nt) database or Refseq, for taxonomic binning of assemblies. Using full alignments captures the synonymous and non-coding variation, as well as the positional relationship of features, without needing to annotate protein coding regions or translate sequences. Several authors have posited that nucleotide alignment lacks sensitivity to classify divergent taxa^6,7^; however, we show this assumption to be false when classifying contigs. Second, our approach leverages the absolute nucleotide identity (ANI) of alignments to collapse taxonomic assignments to higher ranks based on accepted ANI thresholds for defining taxa. Third, we incorporate uncertainty between alignments using a voting algorithm to collapse assignments up the taxonomic tree, and fourth, we adjust for known inaccuracies in reference databases by detecting and correcting misannotated contigs. We show that these advancements, especially when paired with long-read sequencing, enable the automated identification and characterization of microbes in complex microbial communities.

## Results

### Evaluation of taxonomic binning accuracy with mock microbial communities and known organisms

We first evaluate BugSplit using three commonly used benchmarking datasets generated with third-generation sequencers: the ZymoBIOMICS Even and Log datasets are mock microbial communities of eight bacteria and two yeasts, with varying abundance, sequenced on an ONT GridlON^21^, and the ZymoBIOMICS Gut Microbiome Standard containing 19 bacteria (five of which are strains of the same species) and two yeasts sequenced on a PacBio Sequel II. We compare BugSplit with DIAMOND+MEGAN-LR and MMseqs2, two popular tools for taxonomic binning of contigs, and use the official CAMI AMBER tool to assess taxonomic binning performance^6,7,22^. AMBER’s calculation of performance metrics has been previously reported and is summarized in the methods, along with methods for assembly and polishing of each metagenomic community.

On average, BugSplit binned contigs to species bins with an absolute F1-score 33.0% better than DIAMOND+MEGAN-LR and 54.9% better than MMseqs2. These advantages are maintained across all taxonomic ranks, but decrease in magnitude at higher ranks where the performance of DIAMOND+MEGAN-LR and MMseqs2 improves (Figure 1). BugSplit largely demonstrates superior accuracy by improving classification completeness while maintaining or surpassing the classification purity of alternative tools. At the species level, average classification purity was within 6% of DIAMOND+MEGAN-LR and MMseqs2, while average completeness exceeded DIAMOND+MEGAN-LR by 42.3% and MMseqs2 by 58.4%. Similarly, BugSplit was able to classify, to a species level, 36.6% more contigs than DIAMOND+MEGAN-LR and 50.6% more than MMseqs2, while misclassifying 0.5% fewer contigs than DIAMOND+MEGAN-LR and 3.3% more than MMseqs2. Finally, BugSplit achieves these improvements in accuracy without sacrificing execution time: BugSplit executed these processes faster than DIAMOND+MEGAN-LR and MMseqs2 on all three datasets by 100 minutes or more (Supplementary Table 2).

**Figure 1.**
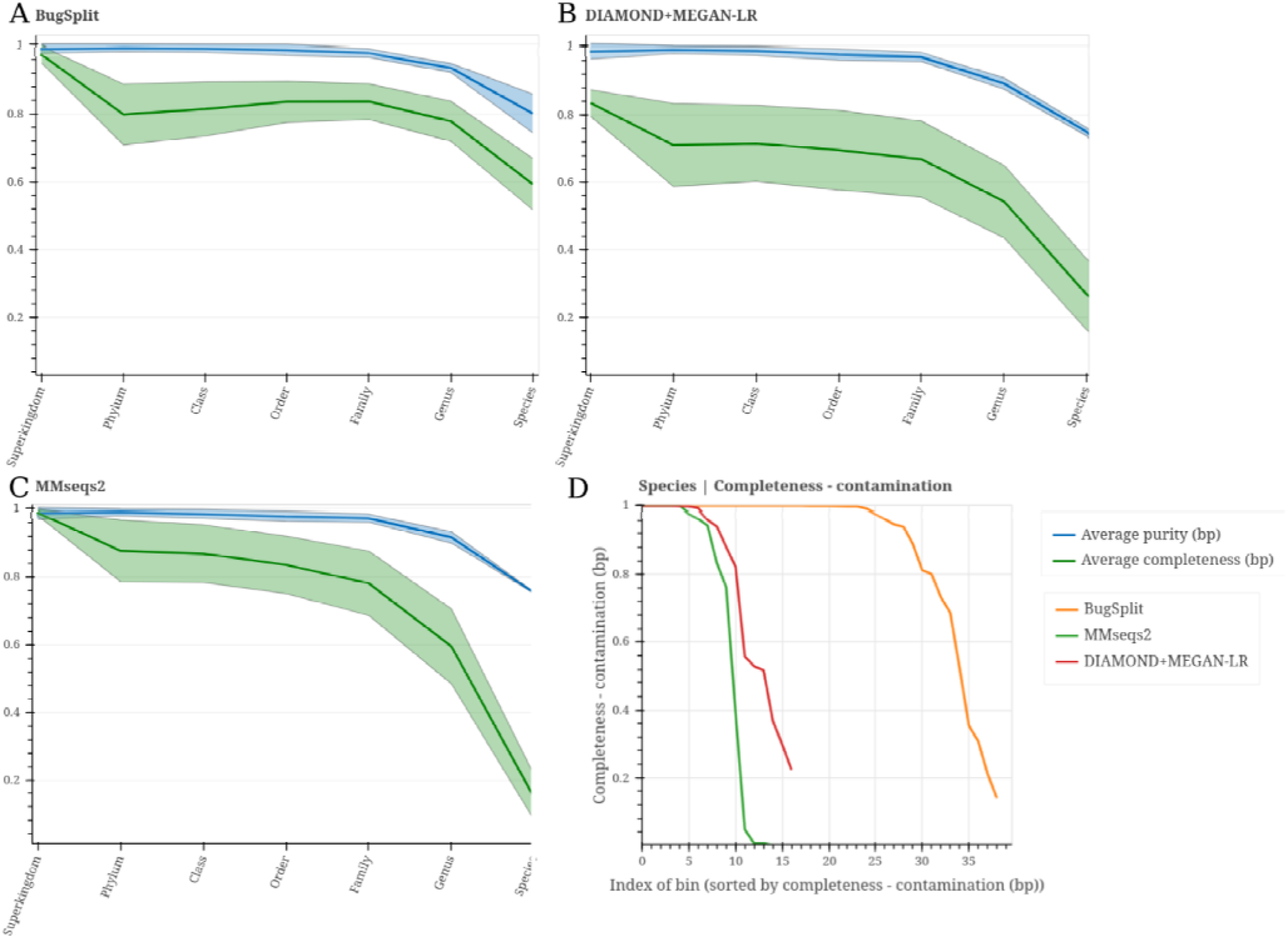
Performance of contig taxonomic classifiers across four datasets: Zymo Even, Zymo Log, Zymo Gut and CAMI High Complexity. Panels A-C: Average bin purity and recall across taxonomic ranks. Shaded bands show the standard error of the metrics. Panel D: BugSplit produces more complete bins with less contamination compared with alternative taxonomic binners.

To highlight the classification performance of BugSplit on a species with close sequence homology yet very different implications for public safety, we apply BugSplit to a case of human anthrax (*Bacillus anthracis* strain Ba0914), sequenced on a MinION at the Centers for Disease Control and Prevention^23^. Our taxonomic binning pipeline classified the complete assembled genome in 1 hour and 20 minutes as *Bacillus anthracis.* MMseqs2 took 3 hours and 48 minutes to identify the bacterial genome as belonging to the family *Bacillaceae,* and DIAMOND+MEGAN-LR took 6 hours 38 minutes to identify the genome as *Bacillus* (most specific taxonomic classification presented for all classifiers).

### Evaluation of taxonomic binning accuracy with novel organisms

To demonstrate that nucleotide alignment retains and even improves on the ability to bin divergent sequences from those in public databases, we apply BugSplit and the other taxonomic binners to two datasets simulating novel organisms. We first analyse the CAMI High-Complexity dataset, a well characterized microbial community comprising 2.80 Gb of contigs from 596 novel organisms generated from a simulated Illumina HiSeq sequencing run, which has been used in other benchmarking studies^2^. We use the CAMI version of Refseq from 2015, predating the publication of organisms in this dataset. BugSplit binned contigs with a genus-level absolute F1-score 16.0% and 15.2% better than DIAMOND+MEGAN-LR and MMseqs2, respectively, which was driven by both superior completeness and purity of bins (7-17.5% better purity, 13.5-14.3% better completeness) (Figure 1). These advantages are maintained at the family rank, where BugSplit exhibited superior purity and completeness by 1 to 3%, respectively.

To demonstrate the impact of BugSplit’s performance on detecting a novel pathogen, we performed nanopore mNGS using sequence-independent single primer amplification of two nasopharyngeal swabs from patients with COVID-19 and from one viral culture of SARS-CoV-2. We generated approximately 5000 to 24,000 reads per sample meeting stringent quality control settings, and assembled 6, 2 and 1 contigs spanning 2,000 to 14,000 base pairs from each sample. We analysed these contigs with BugSplit, DIAMOND+MEGAN-LR and MMseqs2, replacing their databases with an archive of the NCBI nucleotide database from 2019, predating the emergence of SARS-CoV-2.

BugSplit successfully classified all contigs to the genus Betacoronavirus, and correctly classified none of them to the species level. DIAMOND+MEGAN-LR overclassified two contigs, classifying them as Bat SARS-like coronavirus (parent species: Severe acute respiratory syndrome-related coronavirus), and failed to classify the other seven contigs. MMseqs2 classified eight of nine contigs as the less specific Coronaviridae family.

### Improved taxonomic binning enables highly-accurate taxonomic profiling of third-generation metagenomic data

We hypothesized that accurate taxonomic assignment of contigs would improve compositional estimates derived from metagenome sequencing datasets. We have previously shown that alternative long-read taxonomic profilers vastly overestimate the number of species in a sample when applied to mNGS data: in our previous benchmarking study, BugSeq version 1, which was the top performing tool, estimated five times more species than the number of species truly present in a simple mock community (52 vs 10 true species present)^24^. This result was two orders of magnitude better than Centrifuge (>5000 species detected), the tool that currently drives ONT’s official platform EPI2ME^24,25^. As metaFlye^26^ produces length and coverage output of each contig in a metagenomic assembly, we can combine this information with BugSplit taxonomic bin labels to compute the relative abundance of each taxon in a sample in less than five minutes for all third-generation datasets. We additionally include two read-level taxonomic profilers in our comparison: Centrifuge (given its popularity for third-generation metagenomic analysis) and BugSeq version 1 (which has previously been shown to outperform other classifiers including MetaMaps^27^ and CDKAM^28^ on ONT data). Comparison of tools was performed with standardized metrics calculated with OPAL^29^.

Indeed, we find that BugSplit was the top OPAL-ranked among all of the metagenomic profilers assessed, and exhibited the top F1-score (0.81 vs. 0.63 for second-place MMseqs2) when calling species as present or absent across the three benchmark communities (Figure 2). Completeness was also highest after read-based profilers Centrifuge and BugSeq 1; however, Centrifuge called greater than 900 species as present across all datasets (true value: 10-21) and therefore had purity approaching 0, while BugSeq 1 had purity approaching 10%. In comparison, the purity of BugSplit was 89.4%. BugSplit assigned less than 0.02% of bases in each assembly to *Saccharomyces pastorianus,* a hybrid organism of *Saccharomyces cerevisiae* (truly present in each dataset) and *Saccharomyces eubayanus;* filtering these contigs out results in 100% purity for BugSplit across all three long-read datasets. We also find that BugSplit was most accurate identifying the abundance of each species in two of three datasets, and was second to BugSeq 1 for the third dataset, the Zymo Log community (L1 norm error: 0.109 versus 0.073). Organisms that BugSplit failed to detect had the lowest abundance in each dataset, with the exception of *Faecalibacterium prausnitzii*, *Veillonella rogosae* and *Prevotella corporis* from the Zymo Gut dataset, which had their assembled contigs assigned to the genus level. Difficulty in detecting the lowest abundance contigs reflects difficulty in assembling them rather than classifying their contigs; in the Zymo Log dataset, multiple organisms have sequencing coverage below 0.005X and therefore contribute no assembled contigs.

**Figure 2:**
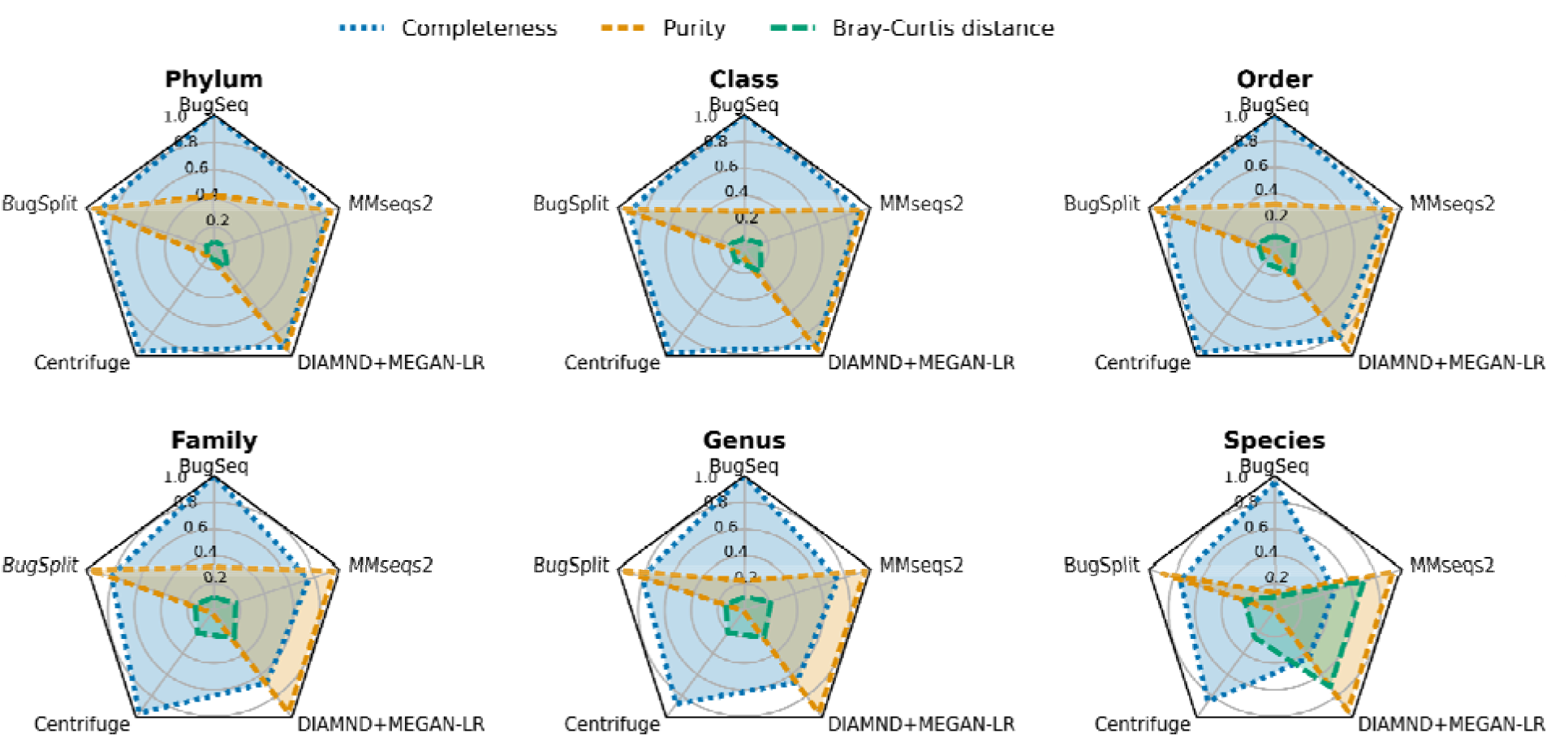
Taxonomic profiling accuracy of five tools across three mock microbial communities sequenced with a long-read sequencer. Greater completeness and purity, and lower Bray-Curtis distance reflect better taxonomic profiling.

**Figure 3:**
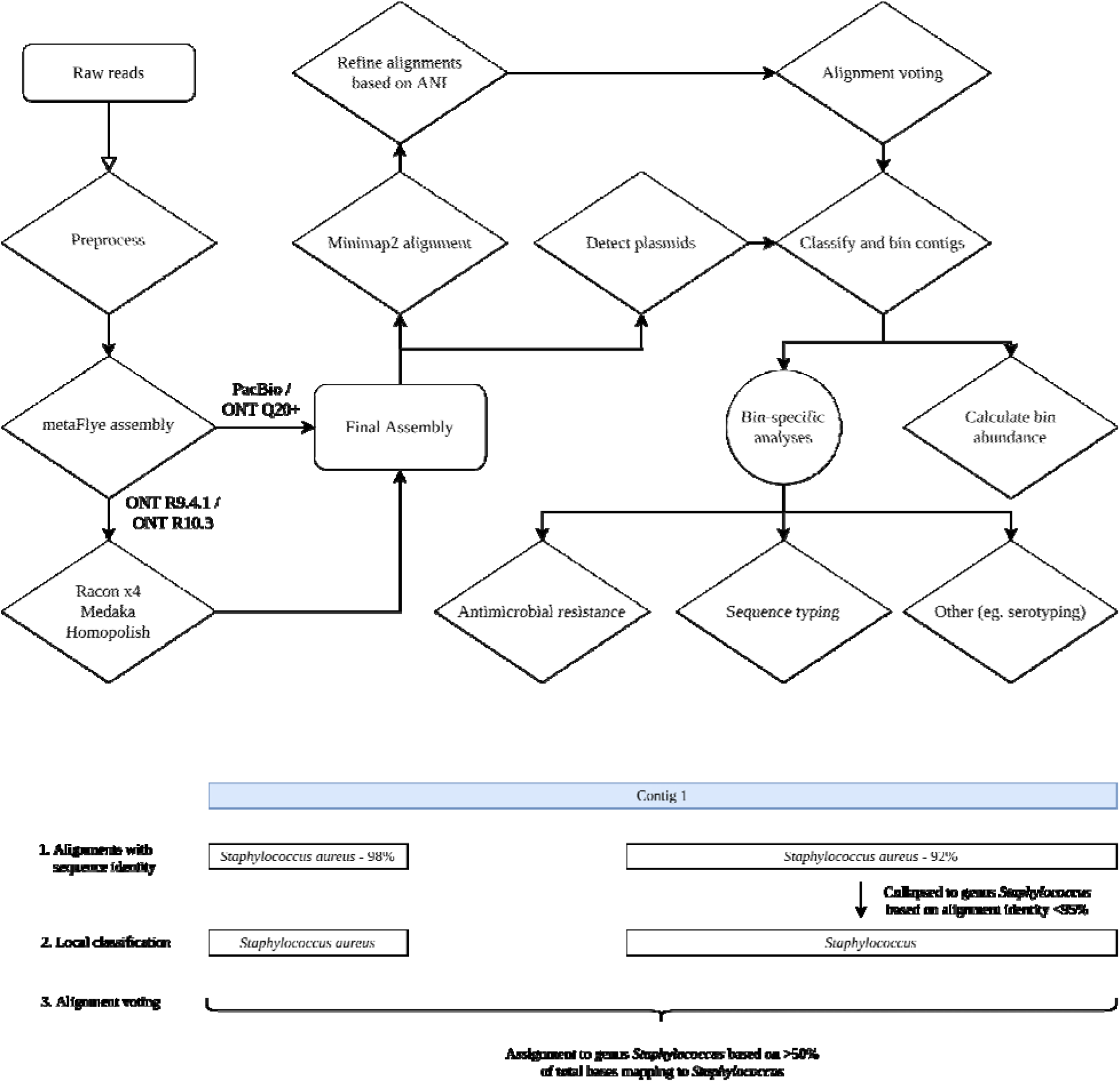
Overview of full BugSplit workflow (top), and example of contig classification algorithm (bottom). Alignments against the reference database are first collapsed up the taxonomic tree based on absolute nucleotide identity. A base-level vote is then performed across all bases of a contig, determining the final taxonomic assignment of the contig based on rankspecific majority thresholds.

### Binning by taxonomic identity enables targeted downstream analysis using tools designed for single-organism whole genome sequencing data

We demonstrate the importance of BugSplit’s species-level taxonomic binning by applying it to a previously reported case of hypervirulent *Klebsiella pneumoniae* liver abscess and its detection using mock blood cultures^30^. Hypervirulence typing of *K. pneumoniae* is important for clinical care, as hypervirulent strains are more likely to cause life-threatening infections and are increasingly resistant to antimicrobials^31^. Two mock blood culture samples from this patient were created by spiking the isolate into blood at 30 CFU/mL, incubating it in an automated blood culture system and sequencing positive blood culture bottles using a nanopore sequencer as previously described^30^. BugSplit successfully detected the only microorganism as *K. pneumoniae* in both samples, and binned sequences belonging to *K. pneumoniae* together, producing bins 97.3% and 94.6% complete with less than 1% contamination using CheckM^32^. In contrast, MMseqs2 produced bins 94.3% and 44.8% complete, and DIAMOND+MEGAN-LR produced bins 29.3 and 25.7% complete. To demonstrate the impact of binning completeness, we implement automatic bioinformatic analyses developed for *K. pneumoniae* whole genome sequencing data (Kleborate^33^) on each bin labelled as *K. pneumoniae*. Kleborate has recently been reported to accurately predict sequence type, serotype and other features of *K. pneumoniae* isolates from WGS data. Kleborate correctly called each BugSplit *K. pneumoniae* bin from both blood culture bottles as sequence type 11 and serotype K47, concordant with bacterial culture and traditional serotyping results. In contrast, Kleborate was unable to determine sequence type for both DIAMOND+MEGAN-LR bins and one of two MMseqs2 bins, and was additionally unable to determine serotype for one of two DIAMOND+MEGAN-LR bins.

By binning at the species level, we can also attribute antimicrobial resistance genes and mutations to specific organisms in a sample. We incorporate ResFinder^34^ into BugSplit and execute it automatically on all taxonomic bins, looking for variants that cause AMR specifically in the taxon assigned to each bin. We additionally implement a modification to ResFinder that automatically adjusts for the higher error rate of nanopore metagenomic assemblies, and corrects indels and erroneous stop codons in resistance conferring loci (see Methods, and publicly available in ResFinder version 4.2). We apply BugSplit and our modified ResFinder tool to a recent dataset of 10 urine samples from patients infected with *Neisseria gonorrhoeae.* Urine underwent mNGS using an ONT GridION, and cultured isolates from the same samples underwent Illumina sequencing for comparison, as previously reported^35^.

BugSplit successfully identified *N. gonorrhoeae* in all 10 samples, ranging from 1.1% to 58.1% of sequenced bases (2.0% to 72.8% of microbial sequenced bases). In contrast to the original Centrifuge-based analysis of this data, and in accordance with the authors’ conclusions that reads classified as *Neisseria lactamica* likely reflect taxonomic misclassification of *N. gonorrhoeae* reads, we find no *N. lactamica* in any of the samples.^35^ Interestingly, we find one sample (original identifier: 301_UB_U) with co-infection with *Ureaplasma urealyticum,* another potential cause of urethritis. The original metagenomic analysis of this data does not discuss this finding, and this sequence was not found in any other samples.

BugSplit successfully assembled and binned together a median of 97.1% (range: 67.0-98.3%) of the *N. gonorrhoeae* genome, as determined by CheckM on the *N. gonorrhoeae* bins, with a maximum of 0.9% contamination. DIAMOND+MEGAN-LR and MMseqs2 binned a median of 11.4% (range: 0% to 26.7%) and 22.4% (range: 0% to 65.5%) of the *N. gonorrhoeae* genome, respectively, with a maximum of 0.2% contamination. When our modified ResFinder is run on the *N. gonorrhoeae* bins created with BugSplit, we identify variants conferring AMR to cephalosporins, quinolones, penicillins, macrolides and tetracyclines with 96.6% sensitivity (28/29 variants detected) and 100% specificity as compared with Illumina sequencing of cultured isolates from the same patients, and the authors’ original pathogen-specific analysis^36^. In comparison, DIAMOND+MEGAN-LR detected 13.8% of variants (4/29) and MMseqs2 detected 24.1% (7/29) variants conferring AMR.

**Table 1.** Antimicrobial resistance prediction of BugSplit, applied to nanopore mNGS of urine, compared with Illumina sequencing, of *Neisseria gonorrhoeae* infections. **Bold** = variant detected by BugSplit, concordant with Illumina sequencing of *N. gonorrhoeae* isolates Red = variant missed by BugSplit, compared with Illumina sequencing of *N. gonorrhoeae* isolates. The mtR gene was not assembled for the single missing variant within this region. N/A = variant position not recovered by BugSplit; no variant present

|  | **23S**<br>**(Macrolides)** |  | **gyrA**<br>**(Quinolones)** |  | **mtrR (Penicillins,**<br>**Tetracyclines,**<br>**Cephalosporins,**<br>**Macrolides)** |  |  | **pilQ**<br>**(Macr**<br>**olides)** | **ponA**<br>**(Penici**<br>**llins)** | **rpsJ**<br>**(Tetra**<br>**cycline**<br>**s)** |
| --- | --- | --- | --- | --- | --- | --- | --- | --- | --- | --- |
| **Sample /**<br>**Resistance**<br>**Variant** | **c.2045**<br>**A>G** | **c.2597**<br>**C>T** | **p.S91F** | **p.D95**<br>**N/G** | **p.A39**<br>**T** | **p.G45**<br>**D** | **Deletio**<br>**n** | **p.E666**<br>**K** | **p.L421**<br>**P** | **p.V57**<br>**M/L/Q** |
| 202 | A | C | S | D | A | G | **deletion** | E | **P** | V |
| 206 | A | C | **F** | A | **T** | N/A | N/A | E | L | **M** |
| 250 | A | C | S | D | A | G | **deletion** | E | **P** | V |
| 271 | A | C | **F** | A | **T** | G | WT | E | L | **M** |
| 294 | A | C | **F** | A | **T** | G | WT | E | L | **M** |
| 301 | A | C | **F** | **G** | A | G | **deletion** | E | **P** | **M** |
| 303 | A | C | **F** | A | **T** | G | WT | E | L | **M** |
| 304 | A | C | S | D | **T** | G | WT | E | L | **M** |
| 314 | A | C | S | D | A | G | **deletion** | E | **P** | V |
| 315 | A | C | **F** | A | **T** | G | WT | E | **P** | **M** |

## Discussion

As mNGS becomes more ubiquitous and is applied to clinical, environmental, biodefense and other samples encompassing incredible microbial diversity, methods to characterize the genome and functional capacity of each organism in a sample become increasingly important. Here, we demonstrate that nucleotide alignment of contigs against a reference database enables significantly improved taxonomic binning of metagenomic assemblies when compared to tools that rely on protein alignments. To evaluate the performance of this approach, we performed taxonomic binning of simulated and real sequencing data from three sequencing technologies and microbial communities containing one to over 500 organisms. BugSplit can classify organisms with available reference genomes to the species level with significantly greater completeness than alternative approaches while preserving bin purity. Our results on simulated novel organisms demonstrate that nucleotide alignment retains sensitivity to classify divergent organisms, with precision to place them on the taxonomic tree at the appropriate rank.

We demonstrate through several use cases how these improvements in taxonomic binning unlock downstream analyses not feasible with current taxonomic binners. Our application of mNGS and BugSplit to the detection of SARS-CoV-2 before any reference sequences were available highlights the power of broadly deploying mNGS with optimal taxonomic binning for pathogen surveillance and pandemic prevention by detecting pathogens earlier. The robust binning provided by BugSplit also allows for automated analysis of taxonomic bins using tools designed for single organism whole genome sequencing data. We demonstrate that it is possible to accurately predict the sequence type, serotype and antimicrobial resistance of organisms directly from clinical samples such as blood cultures and urine. We additionally demonstrate how taxonomic binning may be used to identify unknown organisms of bioterrorism potential, such as *Bacillus anthracis*. As diagnostic microbiology laboratories adopt NGS for bacterial isolate and metagenomic sequencing, automated tools to detect pathogens and leverage the existing body of pathogen-specific bioinformatic analyses will enable a faster, easier transition to NGS. We anticipate that this technology will be broadly useful for the detection and characterization of organisms from diverse samples, and can be greatly expanded upon to support analyses that are application and domain specific.

We anticipate several improvements that will further refine taxonomic bins produced by BugSplit. The most substantial impact is likely to be the inclusion of assembly graph topology into the binning algorithm to improve strain-level resolution. Currently, metagenomic assemblers only output a single contig for conserved regions between different strains of a single species; using BugSplit, these contigs are assigned to the common species and placed in a species-level bin. By incorporating graph topology and linkage of contigs, we will be able to mitigate this limitation and place the contig in multiple strain-level taxonomic bins. Further exploration of the parameter space of BugSplit may also result in improved binning. For example, minimap2 could be tuned for greater alignment recall while preserving precision than its default “map-ont” setting, and voting coverage thresholds may be able to be tuned for improved classification of contigs across the taxonomic hierarchy. Ultimately, we expect to adopt a strategy that will allow optimal values for key parameters to be determined by the taxonomic lineage of alignments.

BugSplit is a highly accurate tool for taxonomic binning and profiling of third-generation metagenomic data with computing speeds faster than comparable workflows. We show that using BugSplit to bin metagenomic assemblies has several substantial downstream effects, including enabling highly similar species discrimination and identification, novel species identification and universal, pathogen-agnostic taxonomic profiling. When combined with automated assembly, polishing and post-processing of bins, we demonstrate that detecting pathogens, strain-typing them and accurately predicting their antimicrobial resistance directly from complex samples with mNGS becomes feasible.

## Methods

### BugSplit Preprocessing

BugSplit uses Nextflow running on AWS Batch to orchestrate processing of sequencing data in the cloud. In brief, nanopore reads undergo demultiplexing and adapter trimming with qcat^37^. Nanopore and PacBio (now added into the pipeline) next undergo quality control with prinseq-lite^38^, filtering reads with a mean Phred score less than 7, a DUST complexity score less than 7 or a read length less than 100 base pairs. Finally, reads are aligned with minimap2^39^ using default parameters against a database containing common non-microbial host genomes, including human, mouse, rat, pig, cow and chicken, to focus assembly on microorganisms. Reads unaligned to host genomes are retained and progress to assembly.

After preprocessing, reads are assembled with metaFlye^26^, preserving strain heterogeneity. Assemblies built from ONT R9.4.1 or R10.3 reads undergo four rounds of Racon^40^ polishing, one round of Medaka^41^ and one round of Homopolish^42^, in accordance with recent assembly benchmarking^43^. A mash database^44^, published by the mash authors and comprising all genomes and plasmid sequences in Refseq (https://gembox.cbcb.umd.edu/mash/refseq.genomes%2Bplasmid.k21s1000.msh) is used for homology search with Homopolish. Racon and Medaka are executed on g4dn-class instances via AWS Batch. PacBio HiFi and ONT Q20+ assemblies do not undergo polishing beyond that included in metaFlye. An entire GridION flowcell using R9.4.1 pores can be assembled and polished to Q40 in less than 6.5 hours, approximately 5 hours faster than using CPUs alone (Supplementary Material).

### Taxonomic Binning of Contigs

Contigs are first aligned to a reference database such as RefSeq or NCBI nucleotide database (nt) with minimap2. We use the default ‘map-ont’ preset of minimap2, as it provides the greatest sensitivity for nucleotide alignment out of all minimap2 presets and performs comparably to nucleotide BLAST^45^. We evaluated replacing minimap2 with an alternative local nucleotide aligner, discontiguous nucleotide megaBLASTN^46^, however this approach was too slow for practical purposes (Supplementary Material). As alignments are made to individual genomes representing a single strain of an organism, the taxonomic identification of each retained alignment is reassigned to internal nodes on the taxonomic tree based on absolute nucleotide identity. Based on the previous identification of ANI thresholds to define a species and genus, as well as the current error rates for metagenomic assembly and the lack of strain representation in public reference databases, we reassign any alignment to the reference database with 95-99% ANI to the species level, with 62-94.9% ANI to the genus level, and with less than 62% ANI to the superkingdom (highest rank before root) level^47–51^. Alignments with greater than 99% ANI are retained at the strain level.

As minimap2 randomly picks a primary alignment if there are multiple alignments with equal top score, we collapse equally good top hits to their lowest common ancestor. Alignments to collapse are identified as secondary alignments with equal dynamic programming score of the max scoring segment in the alignment (“ms” minimap2 SAM tag) to a non-secondary alignment, covering the exact same region of a query contig as the non-secondary alignment.

Next, we implement a voting algorithm to assign contigs to the taxonomic node encompassing a certain percentage of all bases in the contig, aggregating the alignments from above. We again parameterize this vote using accepted definitions of species and previous studies utilizing ANI, requiring 95% and 70% of bases in a contig to map to a strain or species for the contig to be assigned to that strain or species^52–54^, respectively. For ranks above species (eg. genus), we use a majority vote, assigning the contig to the deepest taxon encompassing at least 50% of all bases in a contig, as this approach has previously been reported to perform well^55^. In summary, for a contig to be assigned to a species, it must have at least 70% of its bases with 95% or more ANI mapped to a reference sequence.

NCBI assigns plasmids to the taxonomy identity of their host bacteria in which they were first sequenced^56,57^. This can be misleading due to plasmid conjugation and the ability of plasmids to replicate in organisms across phylogenetic subgroups. We implement a mechanism to recover and correct taxon labels of plasmid sequences. In brief, plasmid sequences are identified with PlasmidFinder^58^, and their taxonomic identities are overridden to that of “plasmid sequences” (NCBI taxon 36549). Full commands and versions for each program are available in the supplementary material.

### Abundance Calculation

The relative abundance of a taxon *t*, in terms of percent of total nucleic acid in a sample, can be approximated using the sequencing depth and length of all contigs *c* assigned to *t* (*c_t_*), divided by the total size of sequencing data:

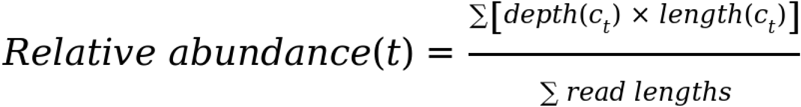

Abundance in bases can be summed up the taxonomic tree to calculate cumulative bases assigned to each taxon, yielding relative abundance at all ranks, in an approach similar to Sczyrba et al^2^.

### Accuracy Assessment and Comparison to Alternative Tools

We use AMBER^22^ version 2.0.2 to assess the performance of each tool binning contigs to taxa. Bin completeness was calculated as the average fraction of true positive base pairs in each predicted bin from the true bin size. Bin purity was calculated as the average fraction of true positive base pairs in each predicted bin. We use OPAL^29^ version 1.0.10 to assess the taxonomic profiling performance of each tool. The default OPAL ranking scheme was used to identify the top taxonomic profiler. OPAL’s purity is calculated as the number of taxa correctly predicted as present in a sample divided by all predicted taxa at that rank. OPAL’s completeness is calculated as the number of taxa correctly predicted as present in a sample divided by all taxa present at that rank. Completeness and purity for both AMBER^22^ and OPAL^29^ range from 0 (worst) to 1 (best). Further calculation details are available in their respective original publications.

MMseqs2 and DIAMOND were run with the NCBI non-redundant amino acid database as suggested by their authors. All databases were downloaded on May 15, 2021, and all tools were run with 96 threads and 768Gb of RAM available to them. The CAMI comparison used the 2015 Refseq database and NCBI taxonomy as provided by the CAMI authors.^2^ The built-in taxonomy of MEGAN-LR was replaced by placing ncbi.tre and ncbi.map, a Newick formatted NCBI taxonomy, in the working directory. These files were generated by converting the NCBI taxonomy files (names.dmp and nodes.dmp) provided with the CAMI datasets into Newick format with the Python taxonomy package^59^.

Ground truth classifications were generated for all datasets except for CAMI, which used the gold standard contig classifications provided by CAMI. Ground truths were generated by comparing each contig in our metagenomic assembly to the reference genome of each organism contained within the mock microbial community using MegaBLASTN. The taxonomic identification of the top BLAST hit for each contig was determined to be its gold standard assignment.

### Application to detection of an emerging coronavirus, hypervirulent *Klebsiella pneumoniae* and *Neisseria gonorrhoeae* infections

For the detection of an emerging coronavirus, nasopharyngeal swabs (n = 2) were collected as part of routine testing at Vancouver General Hospital during Fall 2020 (*ORF1ab* C_t_ values = 14.7, 20.6) and cultured SARS-CoV-2 viral particles (*RdRp* C_t_ value = 18.3) were obtained from the BC Centre for Disease Control Public Health Laboratory. Both clinical samples and cultured virions were extracted and randomly amplified through sequence-independent single-primer amplification as previously described^60^. Samples were sequenced on Oxford Nanopore MinION devices and basecalling was performed with Guppy (Oxford Nanopore Technologies). Ethics approval for collection of nasopharyngeal swabs was obtained from the University of British Columbia (H20-02152).

For the application to human anthrax, hypervirulent *K. pneumoniae*, and *N. gonorrhoeae* infections, raw data was downloaded from the NCBI accessions listed below in Data Availability and submitted to BugSplit. In brief, reads were preprocessed, assembled and polished as detailed above in BugSplit preprocessing. Binning completion and contamination were assessed with CheckM using the default CheckM database. The NCBI nucleotide database from 2019 was downloaded from the second CAMI challenge (https://openstack.cebitec.uni-bielefeld.de:8080/swift/v1/CAMI_2_DATABASES/ncbi_blast/nt.gz) and used in place of BugSplit’s default database for the emerging coronavirus application.

### Modifications to ResFinder to accommodate for insertions, deletions and stop codons in assemblies with high error rates

By default, ResFinder^34^ (which includes the PointFinder^61^ module) performs a BLASTN^46^ alignment of the query assembly against a database of resistance loci. PointFinder scans each alignment and identifies all differences, including insertions and deletions (indels), between the query assembly and reference locus for further annotation. In the event of a stop codon (nucleotides TAG, TAA or TGA) within a locus, PointFinder terminates its search for variants in the region upstream to the stop codon. Full details are available in the original PointFinder^61^ methods. We base our modifications to PointFinder on the previously demonstrated observation that frameshifts and stop codons in third-generation assemblies are more likely to reflect sequencing and assembly errors than true sequence variation^62,63^. We modify PointFinder to not halt its search for variants along a resistance loci if it encounters a stop codon. We additionally modify PointFinder to shift alignments around indels, maintaining the reading frame, in an approach similar to more general frameshift correction tools^62,63^. Our modified PointFinder has been incorporated into ResFinder version 4.2 and can be activated with the ‘-ii’ (Ignore Indels) and ‘-ic’ (Ignore stop Codons) flags.

## Supporting information

AMBER Output File

OPAL Output File

## Data Availability

A public instance of BugSplit is freely available for academic use at https://bugseq.com/academic. Acceptable inputs include FASTQ files from one or more samples sequenced on an Illumina, PacBio or ONT sequencer. Paired Illumina FASTQ files are also accepted. Outputs comprise taxonomic profiling in visual (HTML) and Kraken-report format, taxonomic bins in FASTA format, and additional bin-specific analyses as detailed above in textual and visual formats.

Benchmarking data was downloaded from:

*Bacillus anthracis* whole genome nanopore sequencing: SRA accession SRR10088696 ZymoBIOMICS Even nanopore mNGS: SRA accession ERR3152364

ZymoBIOMICS Log nanopore mNGS: SRA accession ERR3152366

ZymoBIOMICS Gut PacBio HiFi mNGS: SRA accession SRR13128014

CAMI High Complexity gold standard assembly and ground truth labels: https://openstack.cebitec.uni-bielefeld.de:8080/swift/v1/CAMI_I_HIGH using the CAMI downloader.

COVID-19 nanopore mNGS data: Bioproject Accession Number PRJNA766077

Hypervirulent *Klebsiella pneumoniae* nanopore mNGS data: NCBI BioProject PRJNA663005 *Neisseria gonorrhoeae* nanopore mNGS data: NCBI BioProject PRJEB35173

NCBI nt database from 2019: https://openstack.cebitec.uni-bielefeld.de:8080/swift/v1/CAMI_2_DATABASES/ncbi_blast/nt.gz

Our error-tolerant mode has been integrated into ResFinder/PointFinder and is freely available at: https://bitbucket.org/genomicepidemiology/resfinder/src/master/. Error-tolerant mode can be activated with the ‘-ii’ and ‘-ic’ command line flags.

## Notes

### Competing Interest Statement

IC is an employee of BugSeq Bioinformatics Inc. SDC is a shareholder of BugSeq Bioinformatics Inc.

## References

1. Kayani, M. U. R., Huang, W., Feng, R. & Chen, L. Genome-resolved metagenomics using environmental and clinical samples. Brief. Bioinform. 22, (2021).

2. Sczyrba, A. et al. Critical Assessment of Metagenome Interpretation—a benchmark of metagenomics software. Nat. Methods 14, 1063–1071 (2017).

3. Meyer, F. et al. Critical Assessment of Metagenome Interpretation - the second round of challenges. 2021.07.12.451567 https://www.biorxiv.org/content/10.1101/2021.07.12.451567v1 (2021) doi:10.1101/2021.07.12.451567.

4. Breitwieser, F. P., Lu, J. & Salzberg, S. L. A review of methods and databases for metagenomic classification and assembly. Brief. Bioinform. 20, 1125–1136 (2019).

5. Vandenberg, O., Martiny, D., Rochas, O., van Belkum, A. & Kozlakidis, Z. Considerations for diagnostic COVID-19 tests. Nat. Rev. Microbiol. 19, 171–183 (2021).

6. Mirdita, M., Steinegger, M., Breitwieser, F., Söding, J. & Levy Karin, E. Fast and sensitive taxonomic assignment to metagenomic contigs. Bioinformatics 37, 3029–3031 (2021).

7. Huson, D. H. et al. MEGAN-LR: new algorithms allow accurate binning and easy interactive exploration of metagenomic long reads and contigs. Biol. Direct 13, 6 (2018).

8. Bağcı, C., Patz, S. & Huson, D. H. DIAMOND+MEGAN: Fast and Easy Taxonomic and Functional Analysis of Short and Long Microbiome Sequences. Curr. Protoc. 1, e59 (2021).

9. von Meijenfeldt, F. A. B., Arkhipova, K., Cambuy, D. D., Coutinho, F. H. & Dutilh, B. E. Robust taxonomic classification of uncharted microbial sequences and bins with CAT and BAT. Genome Biol. 20, 217 (2019).

10. Gregor, I., Dröge, J., Schirmer, M., Quince, C. & McHardy, A. C. PhyloPythiaS+: a self-training method for the rapid reconstruction of low-ranking taxonomic bins from metagenomes. PeerJ 4, e1603 (2016).

11. Chaumeil, P. A., Mussig, A. J., Hugenholtz, P. & Parks, D. H. GTDB-Tk: a toolkit to classify genomes with the Genome Taxonomy Database. Bioinformatics 36, 1925–1927 (2020).

12. Gehrig, J. L. et al. Finding the right fit: A comprehensive evaluation of short-read and long-read sequencing approaches to maximize the utility of clinical microbiome data. 2021.08.31.458285 https://www.biorxiv.org/content/10.1101/2021.08.31.458285v1 (2021) doi:10.1101/2021.08.31.458285.

13. Malmstrom, R. R. & Eloe-Fadrosh, E. A. Advancing Genome-Resolved Metagenomics beyond the Shotgun. mSystems 4, e00118–19.

14. Payne, A., Holmes, N., Rakyan, V. & Loose, M. BulkVis: a graphical viewer for Oxford nanopore bulk FAST5 files. Bioinformatics 35, 2193–2198 (2019).

15. Lang, D. et al. Comparison of the two up-to-date sequencing technologies for genome assembly: HiFi reads of Pacific Biosciences Sequel II system and ultralong reads of Oxford Nanopore. GigaScience 9, (2020).

16. Lal, A. et al. Improving long-read consensus sequencing accuracy with deep learning. 2021.06.28.450238 https://www.biorxiv.org/content/10.1101/2021.06.28.450238v3 (2021) doi:10.1101/2021.06.28.450238.

17. Wick, R. R., Judd, L. M. & Holt, K. E. Performance of neural network basecalling tools for Oxford Nanopore sequencing. Genome Biol. 20, 129 (2019).

18. Petersen, L. M., Martin, I. W., Moschetti, W. E., Kershaw, C. M. & Tsongalis, G. J. Third-Generation Sequencing in the Clinical Laboratory: Exploring the Advantages and Challenges of Nanopore Sequencing. J. Clin. Microbiol. 58, e01315–19.

19. Maguire, M. et al. Precision long-read metagenomics sequencing for food safety by detection and assembly of Shiga toxin-producing Escherichia coli in irrigation water. PLOS ONE 16, e0245172 (2021).

20. Urban, L. et al. Freshwater monitoring by nanopore sequencing. eLife 10, e61504 (2021).

21. Nicholls, S. M., Quick, J. C., Tang, S. & Loman, N. J. Ultra-deep, long-read nanopore sequencing of mock microbial community standards. GigaScience 8, (2019).

22. Meyer, F. et al. AMBER: Assessment of Metagenome BinnERs. GigaScience 7, (2018).

23. McLaughlin, H. P. et al. Rapid Nanopore Whole-Genome Sequencing for Anthrax Emergency Preparedness. Emerg. Infect. Dis. 26, 358–361 (2020).

24. Fan, J., Huang, S. & Chorlton, S. D. BugSeq: a highly accurate cloud platform for long-read metagenomic analyses. BMC Bioinformatics 22, 160 (2021).

25. Kim, D., Song, L., Breitwieser, F. P. & Salzberg, S. L. Centrifuge: rapid and sensitive classification of metagenomic sequences. Genome Res. 26, 1721–1729 (2016).

26. Kolmogorov, M. et al. metaFlye: scalable long-read metagenome assembly using repeat graphs. Nat. Methods 17, 1103–1110 (2020).

27. Dilthey, A. T., Jain, C., Koren, S. & Phillippy, A. M. Strain-level metagenomic assignment and compositional estimation for long reads with MetaMaps. Nat. Commun. 10, 3066 (2019).

28. Bui, V. K. & Wei, C. CDKAM: a taxonomic classification tool using discriminative k-mers and approximate matching strategies. BMC Bioinformatics 21, 468 (2020).

29. Meyer, F. et al. Assessing taxonomic metagenome profilers with OPAL. Genome Biol. 20, 51 (2019).

30. Zhou, M. et al. Comprehensive Pathogen Identification, Antibiotic Resistance, and Virulence Genes Prediction Directly From Simulated Blood Samples and Positive Blood Cultures by Nanopore Metagenomic Sequencing. Front. Genet. 12, 244 (2021).

31. Russo, T. A. & Marr, C. M. Hypervirulent Klebsiella pneumoniae. Clin. Microbiol. Rev. 32, e00001–19.

32. Parks, D. H., Imelfort, M., Skennerton, C. T., Hugenholtz, P. & Tyson, G. W. CheckM: assessing the quality of microbial genomes recovered from isolates, single cells, and metagenomes. Genome Res. 25, 1043–1055 (2015).

33. Lam, M. M. C. et al. A genomic surveillance framework and genotyping tool for Klebsiella pneumoniae and its related species complex. Nat. Commun. 12, 4188 (2021).

34. Bortolaia, V. et al. ResFinder 4.0 for predictions of phenotypes from genotypes. J. Antimicrob. Chemother. 75, 3491–3500 (2020).

35. Street, T. L. et al. Optimizing DNA Extraction Methods for Nanopore Sequencing of Neisseria gonorrhoeae Directly from Urine Samples. J. Clin. Microbiol. 58, e01822–19.

36. Sanderson, N. D. et al. High precision Neisseria gonorrhoeae variant and antimicrobial resistance calling from metagenomic Nanopore sequencing. Genome Res. 30, 1354–1363 (2020).

37. qcat. (Oxford Nanopore Technologies, 2021).

38. Schmieder, R. & Edwards, R. Quality control and preprocessing of metagenomic datasets. Bioinformatics 27, 863–864 (2011).

39. Li, H. Minimap2: pairwise alignment for nucleotide sequences. Bioinformatics 34, 3094–3100 (2018).

40. Vaser, R., Sović, I., Nagarajan, N. & Šikić, M. Fast and accurate de novo genome assembly from long uncorrected reads. Genome Res. 27, 737–746 (2017).

41. Medaka. (Oxford Nanopore Technologies, 2021).

42. Huang, Y. T., Liu, P. Y. & Shih, P. W. Homopolish: a method for the removal of systematic errors in nanopore sequencing by homologous polishing. Genome Biol. 22, 95 (2021).

43. Latorre-Pérez, A., Villalba-Bermell, P., Pascual, J. & Vilanova, C. Assembly methods for nanopore-based metagenomic sequencing: a comparative study. Sci. Rep. 10, 13588 (2020).

44. Ondov, B. D. et al. Mash: fast genome and metagenome distance estimation using MinHash. Genome Biol. 17, 132 (2016).

45. What parameters best resmble blastn. minimap2 GitHub https://github.com/lh3/minimap2/issues/54 (2017).

46. Morgulis, A. et al. Database indexing for production MegaBLAST searches. Bioinformatics 24, 1757–1764 (2008).

47. Ciufo, S. et al. Using average nucleotide identity to improve taxonomic assignments in prokaryotic genomes at the NCBI. Int. J. Syst. Evol. Microbiol. 68, 2386–2392 (2018).

48. Kim, M., Oh, H. S., Park, S. C. & Chun, J. Towards a taxonomic coherence between average nucleotide identity and 16S rRNA gene sequence similarity for species demarcation of prokaryotes. Int. J. Syst. Evol. Microbiol. 64, 346–351.

49. Richter, M. & Rosselló-Móra, R. Shifting the genomic gold standard for the prokaryotic species definition. Proc. Natl. Acad. Sci. 106, 19126–19131 (2009).

50. Barco, R. A. et al. A Genus Definition for Bacteria and Archaea Based on a Standard Genome Relatedness Index. mBio 11, e02475–19.

51. Federhen, S. et al. Toward richer metadata for microbial sequences: replacing strain-level NCBI taxonomy taxids with BioProject, BioSample and Assembly records. Stand. Genomic Sci. 9, 1275 (2014).

52. Goris, J. et al. DNA–DNA hybridization values and their relationship to whole-genome sequence similarities. Int. J. Syst. Evol. Microbiol. 57, 81–91.

53. Konstantinidis, K. T. & Tiedje, J. M. Genomic insights that advance the species definition for prokaryotes. Proc. Natl. Acad. Sci. 102, 2567–2572 (2005).

54. Konstantinidis, K. T., Ramette, A. & Tiedje, J. M. Toward a More Robust Assessment of Intraspecies Diversity, Using Fewer Genetic Markers. Appl. Environ. Microbiol. 72, 7286–7293 (2006).

55. Hanson, N. W., Konwar, K. M. & Hallam, S. J. LCA*: an entropy-based measure for taxonomic assignment within assembled metagenomes. Bioinformatics 32, 3535–3542 (2016).

56. Robertson, J., Bessonov, K., Schonfeld, J. & Nash, J. H. E. Y. 2020. Universal wholesequence-based plasmid typing and its utility to prediction of host range and epidemiological surveillance. Microb. Genomics 6, e000435.

57. Schoch, C. L. et al. NCBI Taxonomy: a comprehensive update on curation, resources and tools. Database 2020, (2020).

58. Carattoli, A. & Hasman, H. PlasmidFinder and In Silico pMLST: Identification and Typing of Plasmid Replicons in Whole-Genome Sequencing (WGS). Methods Mol. Biol. Clifton NJ 2075, 285–294 (2020).

59. Bovee, R. Taxonomy. (One Codex).

60. Gauthier, N. P. G. et al. Nanopore Metagenomic Sequencing for Detection and Characterization of SARS-CoV-2 in Clinical Samples. 2021.08.13.21261922 https://www.medrxiv.org/content/10.1101/2021.08.13.21261922v1 (2021) doi:10.1101/2021.08.13.21261922.

61. Zankari, E. et al. PointFinder: a novel web tool for WGS-based detection of antimicrobial resistance associated with chromosomal point mutations in bacterial pathogens. J. Antimicrob. Chemother. 72, 2764–2768 (2017).

62. Arumugam, K. et al. Annotated bacterial chromosomes from frame-shift-corrected long-read metagenomic data. Microbiome 7, 61 (2019).

63. Hackl, T. et al. proovframe: frameshift-correction for long-read (meta)genomics. 2021.08.23.457338 https://www.biorxiv.org/content/10.1101/2021.08.23.457338v1 (2021) doi:10.1101/2021.08.23.457338.

