## Supplementary Material for "BugSplit: highly accurate taxonomic binning of metagenomic assemblies enables genome-resolved metagenomics"

Table of Contents

Assembly Pipeline Benchmarking	3

Taxonomic Binning Run Time	4

Evaluation of Alternative Alignment	4

Supplementary Files (Attached)	4

BugSplit Commands and Execution	5

### Assembly Pipeline Benchmarking

|  | ZymoBIOMICS Even GridION (ERR3152364) | | ZymoBIOMICS Log GridION (ERR3152366) | |
| --- | --- | --- | --- | --- |
|  | CPU | GPU | CPU | GPU |
| metaFlye | 3:12 | | 4:05 | |
| Racon x4 | 5:20 | 1:08 | 7:27 | 1:22 |
| Medaka | 1:51 | 0:16 | 0:57 | 0:19 |
| Homopolish | 0:46 | | 0:22 | |
| **Total** | 11:09 | 5:22 | 12:51 | 6:08 |

Supplementary Table 1: Cloud-accelerated assembly and polishing times, comparing with and without a GPU. Racon was run with 48 CPUs, and 4 GPUs (Nvidia T4) for the GPU evaluation. Medaka was run with 64 CPUs, and 1 GPU (Nvidia T4) for the GPU evaluation. Threads were set to 64, and batch size was set to 75.

### Taxonomic Binning Run Time

|  | **Zymo Even (ONT)** | **Zymo Log (ONT)** | **Zymo Gut (PacBio)** | **CAMI High-Complexity (Illumina)** |
| --- | --- | --- | --- | --- |
| **BugSplit** | 136 | 79 | 65 | 208 |
| **DIAMOND+MEGAN-LR** | 1003 | 687 | 550 | 185 |
| **MMseqs2** | 226 | 221 | 109 | 14 |

Supplementary Table 2: Execution time in minutes of BugSplit, DIAMOND+MEGAN-LR and MMseqs2 across four benchmarking datasets. All tools were run on AWS r5a.24xl instances, containing 96 CPUs and 768Gb of RAM.

### Evaluation of Alternative Alignment

We attempted to align the CAMI high complexity assembly against a DUST-masked CAMI Refseq database from 2015 using the following command:

`blastn -task dc-megablast -template_type optimal -template_length 18 -best_hit_overhang 0.1 -best_hit_score_edge 0.1 -num_threads 32 -db dust/refseq_2015_db -out CAMI_blast.txt -evalue 0.01 -db_soft_mask 11 -query CAMI.fna`. Execution was not complete at seven days of runtime, and was therefore terminated.

### Supplementary Files (Attached)

AMBER output - all datasets

OPAL output - all datasets

### BugSplit Commands and Execution

The following commands were used to execute each program:

qcat: `qcat -f INPUT.fastq --detect-middle -b output --trim --filter-barcodes`

prinseq-lite: `prinseq-lite.pl -fastq INPUT.fastq -out_good OUT.fastq -out_bad null -ns_max_p 10 -min_qual_mean 7 -lc_method dust -lc_threshold 7 -min_len 100`

metaFlye: `flye --nano-raw INPUT.fastq --out-dir fly_output --plasmids --meta --keep-haplotypes --trestle`

Racon: `racon -m 8 -x -6 -g -8 -w 500 --include-unpolished -t THREADS reads.fastq overlaps.paf assemfly.fna`

Medaka: `medaka_consensus -i INPUT.fastq -d ASSEMBLY.fna -o output -m MEDAKA_MODEL`

Homopolish: `homopolish polish -a ASSEMBLY.fna -s mash_sketch.msh -m HOMOPOLISH_MODEL -o OUTPUT_DIR`

Minimap2: `minimap2 -a -t THREADS --split-prefix temp nt.mmi contigs.fa`

DIAMOND (v2.0.9) was run with the commands:

`diamond blastx -q ASSEMBLY.fna -d nr.dmnd -o ASSEMBLY.daa -F 15 -f 100 --range-culling --top 10 -p 96 -b10 -c1`

`daa-meganizer -i ASSEMBLY.daa -mdb megan-nucl-Jan201.db --longReads --threads 96 --minSupportPercent 0`

MMseqs2: `mmseqs taxonomy contig nr assignments tmpFolder --tax-lineage 2 --majority 0.5 --vote-mode 1 --lca-mode 3 --orf-filter 1 --threads 96`

AMBER: `amber.py -p 1 -o zymo_even --ncbi_nodes_file nodes.dmp --ncbi_names_file names.dmp -g gold_standard.bbx *.binning`

OPAL: `opal.py -o OUTPUT -g GOLD_STANDARD.profile *.profile`
